# Amyloid-β-induced Alteration of Fast and Localized Calcium Elevations in Cultured Astrocytes

**DOI:** 10.1101/2024.09.23.614604

**Authors:** Kaito Nakata, Joe Sakamoto, Kohei Otomo, Masanao Sato, Hirokazu Ishii, Motosuke Tsutsumi, Ryosuke Enoki, Tomomi Nemoto

## Abstract

Alzheimer’s disease (AD) is a progressive neurodegenerative disorder that causes cognitive decline. Uncovering the mechanisms of neurodegeneration in the early stages is essential to establish a treatment for AD. Recent research has proposed the hypothesis that amyloid-β (Aβ) oligomers elicit an excessive glutamate release from astrocytes toward synapses through intracellular free Ca^2+^ ([Ca^2+^]_*i*_) elevations in astrocytes, finally resulting in neuronal dendritic spine loss. Under physiological conditions, astrocytic [Ca^2+^]_*i*_ elevations range spatially from microdomains to network-wide propagation and temporally from milliseconds to tens of seconds. Astrocytic localized and fast [Ca^2+^]_*i*_ elevations might correlate with glutamate release; however, the Aβ-induced alteration of localized, fast astrocytic [Ca^2+^]_*i*_ elevations remains unexplored.

In this study, we quantitatively investigated the Aβ dimers-induced changes in the spatial and temporal patterns of [Ca^2+^]_*i*_ in a primary culture of astrocytes by two-photon excitation spinning-disk confocal microscopy. The frequency of fast [Ca^2+^]_*i*_ elevations occurring locally in astrocytes (≤0.5 s, ≤35 µm^2^) and [Ca^2+^]_*i*_ event occupancy relative to cell area significantly increased after exposure to Aβ dimers.

The effect of Aβ dimers appeared dose-dependently above 500 nM, and these Aβ dimers-induced [Ca^2+^]_*i*_ elevations were primarily mediated by a metabotropic purinergic receptor (P2Y1 receptor) and Ca^2+^ release from the endoplasmic reticulum. Our findings suggest that the Aβ dimers-induced alterations and hyperactivation of astrocytic [Ca^2+^]_*i*_ is a candidate cellular mechanism in the early stages of AD.

## Introduction

Alzheimer’s disease (AD) is a progressive neurodegenerative disorder marked by behavior and cognitive impairment. The most striking pathological features of AD are lesions known as accumulated amyloid-β (Aβ) protein, a peptide comprising 40–43 amino acids, in the brain (Chen et al., 2017). Aβ aggregates into various types of assemblies, including oligomers, fibrils, and senile plaque. Aβ oligomers have been known to induce neurotoxicity and abnormal intracellular free Ca^2+^ ([Ca^2+^]_*i*_) elevations in astrocytes (Talantova et al., 2013). These [Ca^2+^]_*i*_ elevations might trigger excessive glutamate release from the astrocytes to the surrounding synapses, and excessive glutamate levels may induce dendritic spine loss and finally neurodegeneration (Talantova et al., 2013). Nevertheless, the cellular mechanisms of [Ca^2+^]_*i*_ elevations underlying the exposure to Aβ oligomers are not completely understood.

Astrocytes, a major type of glial cells in the central nervous system, regulate synaptic transmission, remove waste products from the brain, and deliver energy fuels to neurons (Ben Haim and Rowitch, 2016; Ding et al., 2022). Through glutamate uptake from the tripartite synapse consisting of pre-, post-synapse, and perisynaptic astrocytic processes (PAP), astrocytes play a vital role in preventing glutamate excitotoxicity (Rose et al., 2018). On the basis of recent studies, researchers suggest that astrocytes also release glutamate, which is involved in the regulation of neuronal activity (Cuellar-Santoyo et al., 2023). These activities are encoded by [Ca^2+^]_*i*_ elevations in the astrocyte (Bazargani and Attwell, 2016). Spatial and temporal patterns vary from microdomains to the astrocyte network and from sub-second to sub-minute scales under physiological conditions (Semyanov et al., 2020). Recent studies have reported that Aβ oligomers increase [Ca^2+^]_*i*_ elevations at the cell size as well as PAP levels *in vivo*, in slice and in primary cultured astrocytes over relatively long time scales, from several seconds to several minutes (Kelly et al., 2023; Bosson et al., 2017). Nevertheless, the effects of Aβ oligomers on localized and fast [Ca^2+^]_*i*_ elevations have not been investigated.

In this study, we quantitatively analyzed [Ca^2+^]_*i*_ elevations induced by Aβ dimers in primary cultured astrocytes by a high-speed imaging technique, two-photon excitation spinning-disk confocal microscopy (Otomo et al., 2015; Otomo et al., 2020; Kamada et al., 2022). As Aβ dimers have been reported to specifically impair synapse structures (Shankar et al., 2008), we investigated the effects of Aβ dimers on [Ca^2+^]_*i*_ elevations in astrocytes. We found that low doses of Aβ dimers (50, 200 nM) exerted no significant effects on the spatial and temporal patterns of [Ca^2+^]_*i*_ elevations, whereas higher concentrations (>500 nM) of Aβ dimers significantly and dose-dependently increased the frequency of [Ca^2+^]_*i*_ elevations and event occupancy relative to cell area. These [Ca^2+^]_*i*_ elevations were mostly inhibited by antagonists for metabotropic purinergic (P2Y1) receptor and Ca^2+^ release from the endoplasmic reticulum. Our findings suggest that Aβ dimer-induced changes in astrocytic [Ca^2+^]_*i*_ could be one of the key cellular mechanisms in the early stages of AD.

## Materials and Methods

### Animals

C57BL6J/Jms Slc mice (Japan SLC) were housed at 22°C–24°C with a standard 12-h light–dark cycle and *ad libitum* access to water and standard chow. All animal experiments were conducted according to ARRIVE guidelines, and all animal care and experimental procedures were approved by the Institutional Animal Care and Use Committee of the National Institute of Natural Sciences and were performed according to the guidelines of the National Institute for Physiological Science (Approval 23A062).

### Preparation of primary cultured astrocytes and viral transfection

Primary cultures of astrocytes were prepared from the cortex of postnatal 0–3 days C57BL6J/Jms Slc mouse of either sex according to a previously published protocol (Kawano et al., 2017). Astrocytes were plated in a 75-cm^2^ cell culture flask in plating medium composed of Dulbecco’s modified Eagle’s medium with GlutaMAX, pyruvate (35050, GIBCO), supplemented with 10% fetal bovine serum (10437, Invitrogen), 0.1% MITO + Serum Extender (355006, BD Biosciences), and penicillin, and cultured at 37°C in humidified 5% CO_2_ atmosphere for 8–10 days. Growing cells were trypsinized and seeded at 3 × 10^3^ cells per 9.5-mm multi-well glass bottom dish (D141400, Matsunami) coated with fibronectin (F0895, Sigma-Aldrich). Aliquots (1 μL) of adeno-associated virus (AAV) (AAV5-GfaABC1D-cyto-GCaMP6f, addgene) were inoculated onto cultured astrocytes on Days 9–11 of culture. The titer of all AAVs was more than 7.0 × 10^12^ genome copies/mL. The infected astrocytes were cultured for an additional 7–10 days. Since the astrocytes were cultured for a total of 17-22 days prior to the Ca^2+^ imaging in our experimental protocol, the astrocytes were assumed to be confluent and well differentiated. The transfection efficiency of AAV with the GfaABC1D promoter is approximately 50%, indicating that we selected GFAP-positive differentiated astrocytes. The culture medium was changed to artificial cerebrospinal fluid (ACSF) consisting of 146 mM NaCl, 2.5 mM KCl, 1 mM CaCl_2_, 4 mM NaOH, 1 mM MgCl_2_, 20 mM d-glucose, 20 mM sucrose, and 10 mM HEPES for at least 2 h before imaging. The ACSF was adjusted to pH 7.4 and 340 mOsm.

### Preparation of Aβ dimers

Synthetic Aβ (1-40) S26C dimers ([AβS26C]2) were purchased from JPT Peptide Technologies (SP-Ab-24_0.5). They were used for dimerization via a disulfide bond. For experiments, the Aβ dimers were dissolved in dimethylsulfoxide (DMSO) to a concentration of 1 mM and diluted in ACSF. The final DMSO concentration was <0.5% in ACSF.

### High-speed intracellular Ca^2+^ imaging

Astrocytic [Ca^2+^]_*i*_ elevations were visualized using a two-photon excitation spinning-disk confocal microscopy system (Otomo et al., 2015; Otomo et al., 2020) composed of a Ti-Sa laser light source (MaiTai eHP DeepSee, Spectra-Physics), a spinning-disk scanner with 100-µm-wide pinholes aligned on the Nipkow disk (CSU-MPϕ100; Yokogawa Electric) (Shimozawa et al., 2013), an inverted microscope (Ti2-E, Nikon), water immersion objective lens (Plan Apo IR 40X, numerical aperture: 1.15, Nikon), an XY controller (Ti2-S-JS, Nikon), an autofocusing system (Ti2-N-NDM-P, Nikon), a stage incubator (STXG-TIZWX-SET, Tokai Hit), and an EM-CCD camera (iXon Ultra 897, Andor Technology), which were controlled by the NIS-Elements software (Nikon). Time-lapse imaging of astrocytes was conducted over 90 s with 100 ms temporal resolutions, which is a higher frame rate than previous studies of Aβ effects in astrocytes (Shah et al., 2022). The oscillating wavelength of the Ti-Sa laser light was selected as 950-nm optimum for GCaMP6f excitation. Cells were perfused continuously at 37°C with ACSF. Aβ dimers, P2Y1 receptor antagonist (MRS2179) (ab120414, Abcam), and inhibitor of sarco-endoplasmic reticulum Ca^2+^-ATPase (cyclopiazonic acid, CPA) (C1530, Sigma-Aldrich) were added using the perfusion system (flow rate 150 µl/min). Ca^2+^ imaging was performed after complete replacement of the solution in the recording dish. Astrocytes without any [Ca^2+^]_*i*_ changes or those with bright fluorescence due to a sign of cell death were excluded from the analysis. The Ca^2+^ imaging data were obtained from a single GFAP-positive astrocyte per culture dish.

### Quantitative analysis of astrocytic Ca^2+^ signals

Raw images were smoothed with a median filter (radius = 3.0 pixel) using the Fiji software(Schindelin et al., 2012), and the average intensity of the extracellular region was subtracted from the smoothed image as a background fluorescence. The cell boundary was semi-manually determined based on the fluorescence images of GCaMP6f. Briefly, a binary image was generated from the average image using the MinError method for image thresholding. Any areas where the outline could not be accurately reproduced were manually corrected. For the quantitative analysis of [Ca^2+^]_*i*_ elevations, we used the Astrocyte Quantification Analysis (AQuA) program of MATLAB (Mathworks) version developed based on an event-based machine-learning model (Wang et al., 2019). The [Ca^2+^]_*i*_ event occupancy was calculated by measuring the total area of [Ca^2+^]_*i*_ elevations divided by the area within the cell boundary. All AQuA parameters used in this study are presented in Table S1. To generate the two-dimensional map for the initiation point of [Ca^2+^]_i_ elevations in each experiment, the output data of AQuA analysis were used for further additional analysis. The total area of all [Ca^2+^]_i_ elevations throughout the Ca^2+^ imaging was extracted from the AQuA output and mapped as a binary image. The centroid of the earliest regions of each [Ca^2+^]_i_ elevation were calculated as average coordinates. The calculated centroids were mapped as cross-haired markers. These analyses were conducted using a custom MATLAB script.

### Data analysis

Firstly, each dataset was classified into five categories (fast-microdomain, fast-region, slow-microdomain, slow-region, global), then statistical analysis was performed using R (version 4.4.1) and RStudio (version 2024.04.2+764) for each category. Frequencies were analyzed by the generalized linear mixed model (GLMM) using “glmer” function in R package lme4 (version 1.1.35.3). Amplitudes and occupancies were analyzed by the generalized linear model (GLM) using “glm” function in base package (version 4.4.1). In GLM/GLMM analysis, Poisson and Gamma distribution was used for a probability distribution of raw counts of [Ca^2+^]_i_ elevation and, the amplitude and occupancy, respectively. The estimated counts of [Ca^2+^]_i_ elevation was converted to frequency by dividing imaging duration (90 s) in most figures. For the occupancy, area within the cell boundary was used as a log-linear offset. The log link function was used for all GLM/GLMM analyses. The treatments (pretreatment, Aβ dimers administration, washout) and the interaction between the treatments and the groups (different Aβ dimers concentrations, and MRS2179 experiments) are fixed effects throughout GLM/ GLMM analysis, and ID, indicating individual cells, is a random effect for GLMM analysis of frequency and occupancy. To analyze the frequency, amplitude, and occupancy in DMSO and CPA experimental groups that have different data structures from others, Wilcoxon’s signed rank sum test (two-sided for DMSO, one-sided for CPA) was performed. The effect of MRS2179 administration on [Ca^2+^]_i_ elevation was analyzed by two-sided Wilcoxon-Mann-Whitney test. All non-parametric tests were performed by using coin package (version 1.4.3). For cells with no [Ca^2+^]_i_ elevation, the amplitudes were missing value (filled with N.A. in dataset), and total area of [Ca^2+^]_i_ elevations were zero. If the amplitude was N.A. for any one of the three treatments, cells were excluded to properly analyze the amplitudes. Because the frequency of global [Ca^2+^]_i_ elevations was zero in most of the cells, the amplitude of global [Ca^2+^]_i_ elevations cannot be analyzed statistically. Plots were created by using ggplot2 (version 3.5.1) and graphics (version 4.4.1) package on R and Excel (Microsoft).

## Results

### Quantification analysis of spontaneous [Ca^2+^]_*i*_ elevations in cultured astrocytes

To visualize astrocytic [Ca^2+^]_*i*_ elevations with high spatial and temporal resolution, we used the high-speed low-invasive two-photon excitation spinning-disk confocal microscopy (Otomo et al., 2015; Otomo et al., 2020) and expressed the genetically-encoded Ca^2+^ probe, GCaMP6f, in the cytosol of primary cortical cultured astrocytes. Diverse spontaneous [Ca^2+^]_*i*_ elevations were observed in different regions of astrocytes, as illustrated in Fig.1A. For the quantitative analysis of [Ca^2+^]_*i*_ elevations, we used an event-based machine-learning model, the AQuA algorithm (Wang et al., 2019) (Fig. 1B). The key parameters that define [Ca^2+^]_*i*_ elevations, such as amplitude (*Δ*F/F_0_), duration time (full-width half-maximum), area, and frequency, were extracted using the AQuA algorithm. The histograms of duration and area revealed that the [Ca^2+^]_*i*_ elevations in astrocytes exhibited a continuous rather than a bimodal distribution (Figs. 1C, D). Therefore, we adopted a criterion in which 50% of the total [Ca^2+^]_*i*_ elevations defined the condition as fast and microdomain. The AQuA analysis for spontaneous astrocytic [Ca^2+^]_*i*_ elevations revealed that 50% of the astrocytic [Ca^2+^]_*i*_ elevations had a duration time of 0.5 s, and the area of that 50% of the [Ca^2+^]_*i*_ elevations was <35 µm^2^. The area of [Ca^2+^]_*i*_ elevations above 400 µm^2^ was associated with Ca^2+^ propagation in astrocytes. On the basis of these results, we classified the patterns of [Ca^2+^]_*i*_ elevations into five categories as follows: “fast-microdomain” (≤0.5 s, ≤35 µm^2^), “slow-microdomain” (>0.5 s, ≤35 µm^2^), “fast-region” (≤0.5 s, >35 µm^2^ and ≤400 µm^2^), “slow-region” (>0.5 s, >35 µm^2^ and ≤400 µm^2^), and “global” (>400 µm^2^) (Fig. 1E). The definition of fast-microdomain was generally comparable to a previous study (Santello et al., 2011).

**Figure 1.**
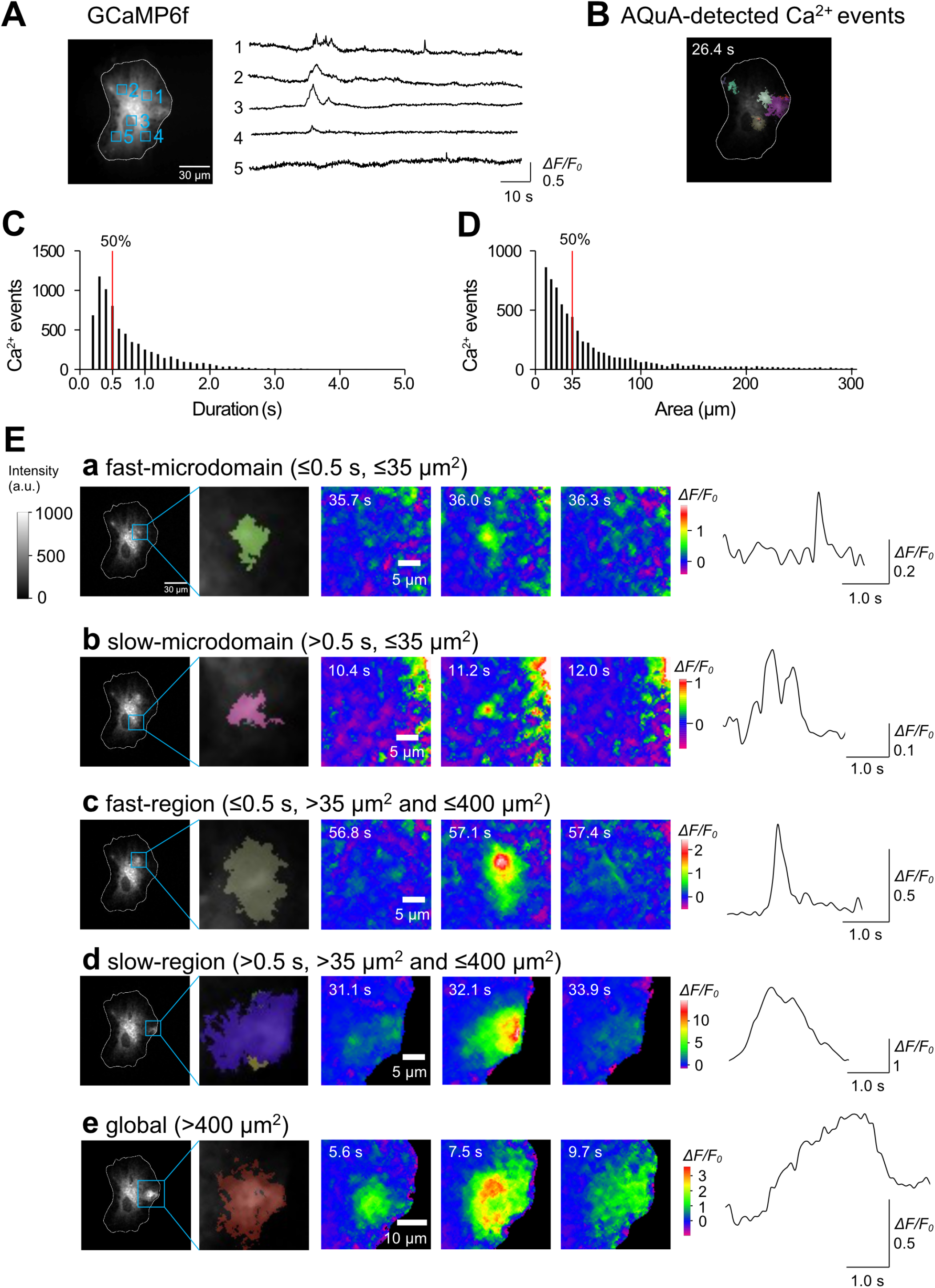
Classification of spatiotemporal [Ca^2+^]_*i*_ patterns in astrocytes. **(A)** Average time projection image (left) and [Ca^2+^]_*i*_ traces (right) of a representative astrocyte. The cell border is indicated by a white line. Boxed regions of interest (ROIs) correspond to the traces on the right. **(B)** Snapshot images of AQuA-detected [Ca^2+^]_*i*_ elevations. Each color in the bottom images represents individual AQuA-detected events. Colors are chosen randomly. **(C)** Histogram of duration time (s). **(D)** Histogram of area (µm^2^). The red vertical line in (C) and (D) indicates 50% of total [Ca^2+^]_*i*_ elevations. **(E)** Representative images and traces of five categories of [Ca^2+^]_*i*_ elevations (a-e). The two left panels show AQuA-detected individual [Ca^2+^]_*i*_ elevations and their magnified images. The three pseudocolor images on the right correspond to the ROIs in the left. Traces of [Ca^2+^]_*i*_ elevations are illustrated on the right. The time is shown in the upper left corner of each pseudocolor image.

### Effect of Aβ dimers on [Ca^2+^]_*i*_ elevations in astrocytes

To ensure the stability and continuity of Ca^2+^ recordings under our experimental conditions, we administered twice the highest concentration of organic solvent (1% DMSO) used in this study and monitored spontaneous [Ca^2+^]_*i*_ elevations (Fig. S1A). We compared the [Ca^2+^]_*i*_ event occupancy relative to cell area during pretreatment and DMSO administration and observed no significant differences (Figs. S1B and C). We also compared the frequency and amplitude of all five categories of [Ca^2+^]_*i*_ elevations (fast-microdomain, slow-microdomain, fast-region, slow-region, and global). Although the sites of [Ca^2+^]_*i*_ elevations in astrocytes changed from time to time during the recording, no significant differences were observed in any categories of [Ca^2+^]_*i*_ elevations (Figs. S1DEF).

Next, we systematically administered different concentrations of Aβ dimers (50 nM, 200 nM, 500 nM, 2 µM, and 5 µM) and analyzed the event occupancy, frequency, and amplitude in all five categories. Representative examples and a statistical comparison are illustrated in Fig. 2, and the results of individual experiments are depicted in Supplemental Figures (Figs. S2–S6).

**Figure 2.**
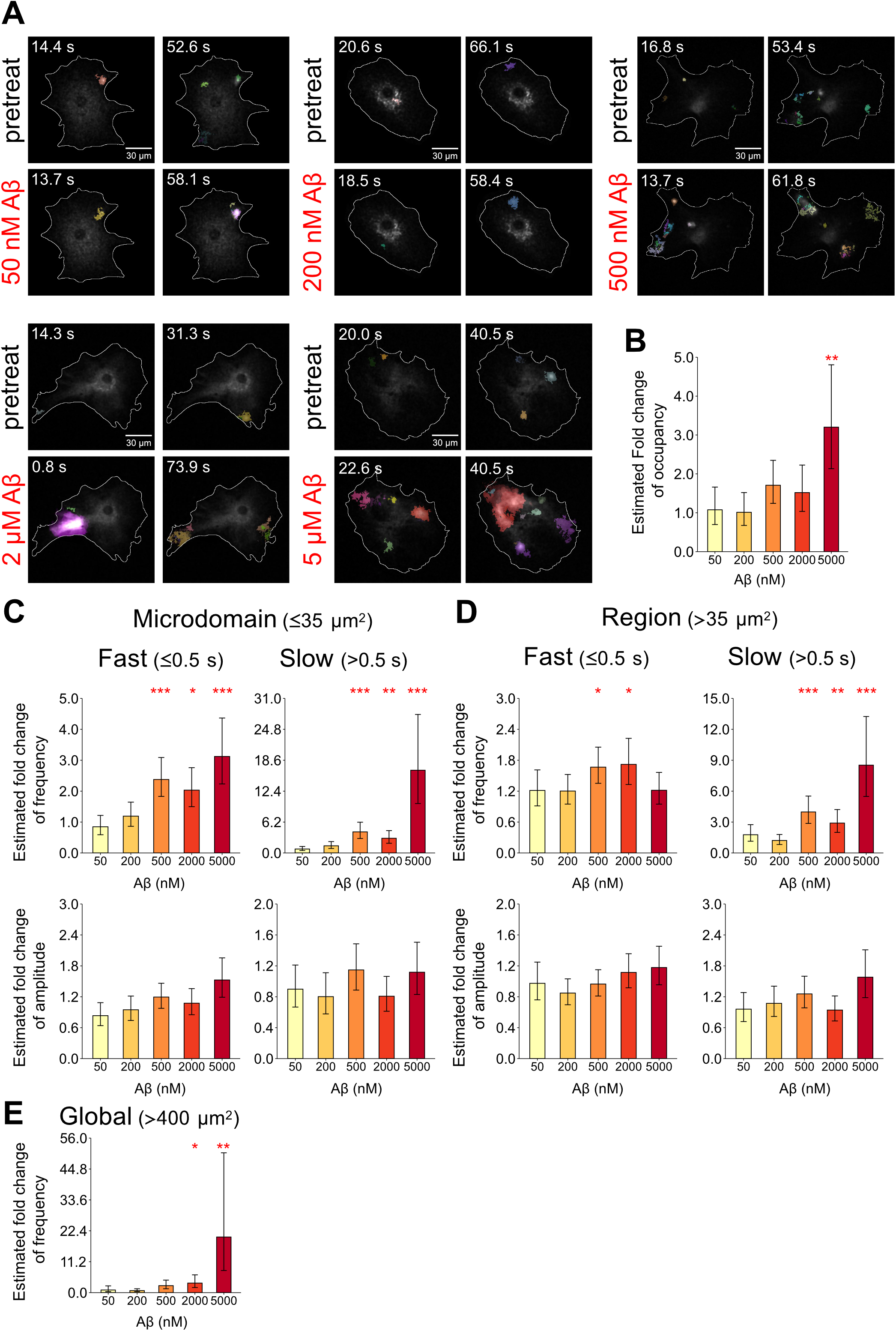
Dose-dependent effects of Aβ dimers on astrocytic [Ca^2+^]_i_ elevations. **(A)** Representative images of astrocytic [Ca^2+^]_i_ elevations displayed with AQuA-detected events (indicated by colored areas). The cell boundary is indicated by a white line. Top: Pretreatment images, bottom: Aβ dimer application images (50 nM, 200 nM, 500 nM, 2 µM, and 5 µM). The time is shown in the upper left corner of each image. The total acquisition time is 90 s. **(B)** Estimated fold change of the event occupancy of [Ca^2+^]_*i*_ elevations under the different Aβ dimer application. **(C)** Dose-dependent changes in the effects of Aβ dimers on the frequency (top) and amplitude (bottom) of microdomain [Ca^2+^]_*i*_ elevations. The categories of fast and slow [Ca^2+^]_*i*_ elevations are depicted in the left and right row graphs. **(D)** Dose-dependent changes in the effects of Aβ dimers on the frequency (top) and amplitude (bottom) of region [Ca^2+^]_*i*_ elevations. The categories of fast and slow [Ca^2+^]_*i*_ elevations are depicted in the left and right row graphs. (E) Dose-dependent changes in the effects of Aβ dimers on the frequency of global [Ca^2+^]_*i*_ elevations. Amplitudes of global [Ca^2+^]_*i*_ elevations were excluded from statistical analyses due to the groups with the small cell number less than 3. In the graphs, individual data (dots) and estimates ± standard error are shown. GLM or GLMM was used to evaluate the difference. *: p < 0.05, **: p < 0.01 (50 nM: n = 7 cells, 200 nM: n = 8 cells, 500 nM: n = 13 cells, 2 µM: n = 9 cells, 5 µM: n = 8 cells). Individual data and statistical results in the graphs are summarized in Supplemental Data Sheet 1.

Low concentrations (50 nM, 200 nM) of Aβ dimers exerted no significant effect on the [Ca^2+^]_*i*_ event occupancy (Figs. S2BC, S3BC) as well as on the frequency and amplitude in all five categories (Figs. S2DEF, S3DEF). We concluded that <200 nM Aβ dimers exerted no detectable effect on spontaneous [Ca^2+^]_*i*_ elevations.

We further explored the effect of higher concentrations of Aβ dimers on [Ca^2+^]_i_ elevations. At 500 nM, 2 µM, and 5 µM concentrations (Figs. S4–S6, Supplementary Movies 1-3), the frequency of fast-microdomain, slow-microdomain, slow-region [Ca^2+^]_*i*_ elevations were significantly increased (Figs. S4DEF, S5DEF, S6DEF). At 500 nM and 2 µM concentrations (Figs. S5–S6), the frequency of fast-region [Ca^2+^]_*i*_ elevations was also significantly increased (Figs. S5F, S6F). At 2 µM and 5 µM concentrations, the frequency of global [Ca^2+^]_*i*_ elevations was significantly increased (Fig. S6E), suggesting that the fast [Ca^2+^]_*i*_ elevations were accumulated and produced a large and long-lasting [Ca^2+^]_*i*_ elevation that spread widely in the astrocyte. At 5 µM concentrations (Fig. S6), the [Ca^2+^]_*i*_ event occupancy were significantly increased. At all concentrations, the amplitude of fast-mocrodomain, slow-microdomain, fast-region, slow-region [Ca^2+^]_*i*_ elevations was not significantly increased. At high concentrations (500 nM, 2 µM, 5 µM), the effects on [Ca^2+^]_*i*_ elevations were not reversible after washout (Figs. S5, 6D, E), suggesting that Aβ dimers were tightly bound to the target molecules or irreversibly altered [Ca^2+^] signaling in astrocytes.

The summary graphs in Fig. 2 show that Aβ dimers exerted dose-dependent effects on the event occupancy, frequency of [Ca^2+^]_*i*_ elevations. The effects of Aβ dimers on [Ca^2+^]_*i*_ elevations were not evident at 200 nM but became significant at >500 nM, indicating a threshold-like response of Aβ dimers on astrocytes (Fig. 2B). In all of the categories, the frequency, but not the amplitude, increased significantly with the concentration of Aβ dimers (Figs. 2C and D). These findings indicate that Aβ dimers dose-dependently increased the [Ca^2+^]_*i*_ elevations in astrocytes. The data for all figures and statistical results are summarized in Supplemental Data Sheet 1.

### Aβ dimers-induced [Ca^2+^]_*i*_ elevations are mediated by P2Y1 receptor activation

We next investigated the mechanism of Aβ dimers-induced [Ca^2+^]_i_ elevations. Previous studies have demonstrated that, under physiological conditions, fast-microdomain [Ca^2+^]_*i*_ elevations involve the activation of purinergic G protein-coupled P2Y1 receptors (Santello et al., 2011; Shigetomi et al., 2018). To examine whether Aβ dimers-induced [Ca^2+^]_*i*_ elevations are mediated by P2Y1 receptors, we analyzed [Ca^2+^]_*i*_ elevations in the presence of the P2Y1 receptor antagonist MRS2179 (20 µM) at an effective concentration of Aβ dimers of 500 nM (Fig. 3A). We observed that the event occupancy, frequency, and amplitude of [Ca^2+^]_*i*_ elevations were unaffected by Aβ dimer administration in the presence of MRS2179 in most categories except fast-region [Ca^2+^]_*i*_ elevations (Figs. 3B–E). MRS2179 administration alone had no significant effect on the [Ca^2+^]_*i*_ elevations (Fig.S7A-D). These findings suggest that Aβ dimers-induced [Ca^2+^]_*i*_ elevations are mostly mediated by signaling involving the activation of the P2Y1 receptor.

**Figure 3.**
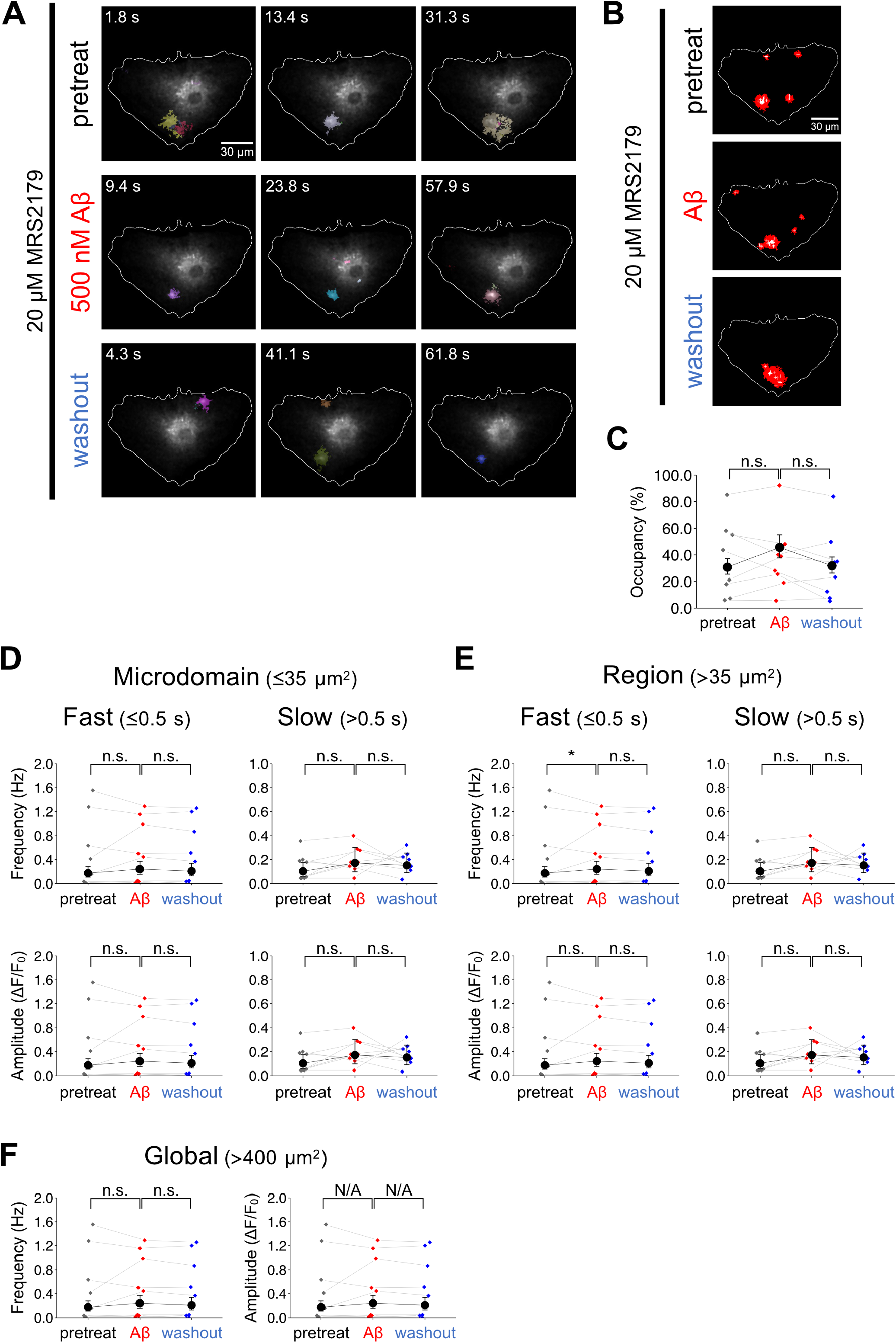
P2Y1 receptor antagonist inhibits Aβ dimers-induced [Ca^2+^]_*i*_ elevations. **(A)** Representative images of astrocytic [Ca^2+^]_*i*_ signals shown with AQuA-detected events. Images of pretreat (top), under 500 µM Aβ dimer application (middle), and after washout (bottom) are shown. Throughout the acquisition, 20 µM MRS2179 was present. The time is shown in the upper left corner of each image. The total acquisition time is 90 s for each condition. **(B)** Comparison of the area of [Ca^2+^]_*i*_ elevations in each condition. The three images represent the event occupancy of [Ca^2+^]_*i*_ elevations in pretreat, Aβ dimer application, and washout. The red area and the white cross indicate the area of [Ca^2+^]_*i*_ elevation and the center of gravity of individual [Ca^2+^]_*i*_ elevations, respectively. **(C)** Comparison of the event occupancy of [Ca^2+^]_*i*_ elevations between pretreat, Aβ dimer application, and washout. **(D, E, F)** Comparison of frequencies (Hz) and amplitudes (*Δ*F/F, %) of [Ca^2+^]_*i*_ elevations between pretreat, Aβ dimer application, and washout in the presence of MRS2179. Data are expressed as estimate ± standard error. GLM or GLMM was used to compare differences (n = 8). Individual data and statistical results in the graphs are summarized in Supplemental Data Sheet 1. n.s.: not significant, N/A: not applicable.

### Aβ dimers-induced [Ca^2+^]_*i*_ elevations are mediated by Ca^2^ release from endoplasmic reticulum

The activation of P2Y1R produces inositol 1,4,5-trisphosphate (IP_3_), which triggers Ca^2+^ release from the endoplasmic reticulum (ER). To further investigate signaling mechanism of Aβ dimers-induced [Ca^2+^]_*i*_ elevations, we analyzed [Ca^2+^]_*i*_ elevations in the presence of cyclopiazonic acid (CPA) (10 µM), a specific inhibitor of sarco-endoplasmic reticulum Ca^2+^-ATPase (SERCA), at an effective concentration of Aβ dimers of 500 nM (Fig. 4). We found that the event occupancy, frequency, and amplitude of [Ca^2+^]_*i*_ elevations were unaffected by Aβ dimer administration in the presence of CPA (Figs. 4B–F). These results indicate that Aβ dimers-induced [Ca^2+^]_*i*_ elevations are mainly mediated by Ca^2+^ release from ER. Notably, [Ca^2+^]_i_ elevations remained detectable in the presence of Aβ dimers and CPA. Since CPA application did not affect the frequency of slow-microdomain [Ca^2+^]_*i*_ elevations (Fig. 4D), it is likely that the slow-microdomain [Ca^2+^]_*i*_ elevation originated from the extracellular space but not from the ER.

**Figure 4.**
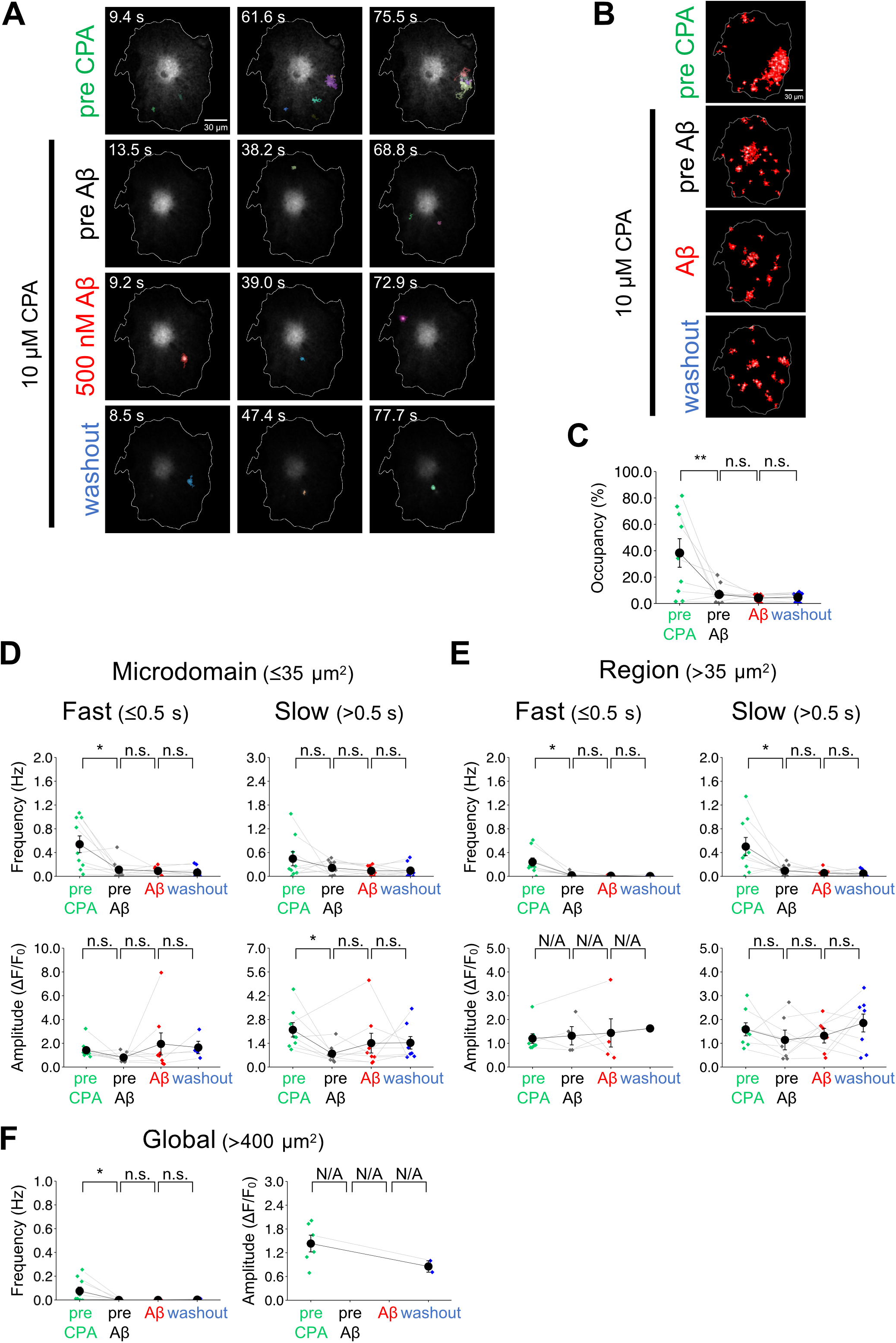
SERCA inhibitor suppressed Aβ dimers-induced [Ca^2+^]_*i*_ elevations. **(A)** Representative images of astrocytic [Ca^2+^]_*i*_ signals shown with AQuA-detected events. Images of pretreat CPA (pre CPA) (top), pretreat Aβ under 10 µM CPA application (pre Aβ) (upper middle), under 500 nM Aβ dimers in the presence of 10 µM CPA application (lower middle), and washout (bottom) in the presence of CPA application are shown. The time is shown in the upper left corner of each image. The total acquisition time is 90 s for each condition. **(B)** Comparison of the area of [Ca^2+^]_*i*_ elevations in each condition. The four images represent the event occupancy of [Ca^2+^]_*i*_ elevations in pre CPA, pre Aβ, Aβ dimer application, and washout. The red area and the white cross indicate the area of [Ca^2+^]_*i*_ elevation and the center of gravity of individual [Ca^2+^]_*i*_ elevations, respectively. **(C)** Comparison of the event occupancy of [Ca^2+^]_*i*_ elevations between pre CPA, pre Aβ, Aβ dimer application, and washout. **(D, E, F)** Comparison of frequencies (Hz) and amplitudes (*Δ*F/F, %) of [Ca^2+^]_*i*_ elevations between pretreat, Aβ dimer application, and washout in the presence of MRS2179. Data are expressed as mean ± s.e.m. One-sided Wilcoxon’s signed rank sum test was used to compare differences (n = 9). Amplitude of global [Ca^2+^]_*i*_ elevation cannot be analyzed statistically as same reason as Fig. 2. Individual data and statistical results in the graphs are summarized in Supplemental Data Sheet 1. n.s.: not significant, N/A: not applicable.

## Discussion

### Aβ dimers-induced alteration of fast-microdomain [Ca^2+^]_*i*_ elevations in astrocytes

We visualized Aβ dimers-induced alteration of [Ca^2+^]_*i*_ elevations in primary cultured astrocytes with high spatial and temporal resolution. We found that >70% of the spontaneous [Ca^2+^]_*i*_ elevations in the astrocytes had a duration of <1 s. Because the speed of Ca^2+^ imaging in previous studies was <1 Hz (Talantova et al., 2013; Pham et al., 2021; Shah et al., 2022), it is possible that most of the [Ca^2+^]_*i*_ elevations were below detectable levels. Notably, we observed that global [Ca^2+^]_*i*_ elevations often accompanied by the Ca^2+^ microdomains (see Supplementary Movies). This result suggests that the continuously occurring microdomain [Ca^2+^]_*i*_ elevation may be the cause of the global [Ca^2+^]_*i*_ elevation.

It has been reported that Ca^2+^ microdomains in astrocytes reflect neuronal activity (Di Castro et al., 2011; Bindocci et al., 2017). Under experimental conditions where both neurons and astrocytes are present and communicate with each other, such as in brain slices or *in vivo*, it is difficult to distinguish which cell type the drug is acting on. We used primary cultured astrocytes as a simple model and analyzed the effects of Aβ dimers without considering the interaction with neurons. To the best of our knowledge, this is the first demonstration of the quantitative analysis of Aβ dimers acting directly on fast [Ca^2+^]_i_ elevations in astrocytes.

### Mechanistic insight into the site of action of Aβ dimers in the astrocyte

We found that Aβ dimers at >500 nM increased the frequency of [Ca^2+^]_*i*_ elevations in astrocytes. Although the mechanism and target site of Aβ dimers remain to be elucidated, our findings suggest that the number of Ca^2+^ channels activated by Aβ dimers is increased in the astrocyte. Under physiological conditions, elevated Ca^2+^ is either taken up into the endoplasmic reticulum (ER) via the sarcoplasmic reticulum Ca^2+^-ATPase (SERCA) or pumped out of the cells via the sodium–calcium exchanger (NCX) on the plasma membrane. Our results suggest that the balance of Ca^2+^ influx and efflux is altered by Aβ dimers.

Previous studies have reported that Aβ oligomers-induced [Ca^2+^]_*i*_ elevations were mediated via Ca^2+^ channel on cell membrane such as transient receptor potential A1 (TRPA1) channels (Paumier et al., 2022) and α7 nicotinic acetylcholine receptors (α7nAchR) (Talantova et al., 2013), and IP_3_ receptors on ER membrane (Alberdi et al., 2013). Importantly, we found that Aβ dimers-induced [Ca^2+^]_*i*_ elevations were inhibited by a P2Y1 receptor antagonist (Fig. 3) and CPA (Fig. 4). Hence, Aβ dimers-induced [Ca^2+^]_*i*_ elevations can be attributed to IP_3_-induced Ca^2+^ release (IICR). A previous study using HeLa cells reported that Ca^2+^ microdomains coincide with the localization of IP_3_ receptors (Thillaiappan et al., 2017). The astrocyte expresses all IP_3_R subtypes in the soma and its processes (Sherwood et al., 2021). The increased event occupancy in the astrocyte (Fig. 2B) could be due to an increased number of IP_3_ receptor sites activated by Aβ dimers. We found that the majority of spontaneous [Ca^2+^]_*i*_ elevations was found to be Ca^2+^ release from the ER into the cytosol via IP_3_ receptors and Aβ dimers-induced [Ca^2+^]_*i*_ elevations were inhibited by CPA (Fig. 4). Hence, Aβ dimers-induced [Ca^2+^]_*i*_ elevations can be primarily attributed to release from the ER. We observed that Aβ dimers-induced fast-region [Ca^2+^]_*i*_ elevations were inhibited by CPA, but not by P2Y1 receptor antagonist. These results indicate Aβ dimers-induced fast-region [Ca^2+^]_*i*_ elevations were mediated by Ca^2+^ release from the ER via other receptors such as metabotropic glutamate receptors (mGluR) or Ca^2+^ influx from extracellular space (Liu et al., 2021). Additionally, we observed the remaining slow-microdomain [Ca^2+^]_*i*_ elevations in the presence of CPA and Aβ dimers, suggesting an extracellular origin. It has been reported that Aβ oligomer-induced [Ca^2+^]_i_ elevations are mediated through the extracellular space via the TRPA1 channel or α7nAchR (Paumier et al., 2022, Talantova et al., 2013). Previous studies have used relatively high molecular weight Aβ (∼ 50 kDa), which may be the reason for the difference in results. Indeed, it has been reported in neuron that Aβ interacts with different receptors depending on its molecular weight (Jarosz-Griffiths et al., 2016).

Identifying the key molecules involved in Aβ oligomer binding will be critical for drug development for AD treatment. Previous studies have demonstrated that various receptor proteins selectively bind to distinct types of Aβ oligomer species (Jarosz-Griffiths et al., 2016). In neurons, Aβ dimers bind β2 adrenergic receptors (Wang et al., 2010). Whether Aβ oligomers bind directly to P2Y1 receptors or bind to another molecule to activate P2Y1R remains incompletely understood. It has been demonstrated that ATP release from CX hemichannels, a major pathway for ATP release from astrocytes, activates astrocyte P2Y1 receptors and induces [Ca^2+^]_*i*_ elevations (Torres et al., 2012). Aβ dimers may initiate ATP release from CX hemichannels and subsequently activate astrocytic P2Y1 receptors.

A recent study identified five morphologically and physiologically distinct astrocyte subtypes in the cerebral cortex and hippocampus of adult mice (Batiuk et al., 2020) and found that the frequency and amplitude of spontaneous astrocyte [Ca^2+^]_i_ elevations differed between layer 1, layers 3-5 of the cortex, and the hippocampus. In our study, the frequency and amplitude of spontaneous [Ca^2+^]_i_ elevations varied from cell to cell. The cell-to-cell variability of these [Ca^2+^]_i_ elevations in this study may be due to the heterogeneity of astrocytes.

### Future perspective

Large-scale drug screening will be required to identify the mechanism of action through which Aβ oligomers activate receptors. The primary culture of astrocytes used in this study allows evaluating the effects of drugs without considering the neuronal inputs. On the other hand, astrocytes have fine structure such as PAPs *in vivo* and in slice. PAPs contact with both pre- and post-synapse and play a role in facilitating neurotransmission. In particular, fast [Ca^2+^]_*i*_ elevations in PAP induce neurotransmitter release such as glutamate, and modulate synapse plasticity (Santello et al., 2011). To elucidate the mechanism of dendritic spine loss in AD, it is crucial to clarify whether Aβ dimer actually alters fast-microdomain [Ca^2+^]_*i*_ elevations in PAPs.

Under physiological conditions, fast-microdomain [Ca^2+^]_*i*_ elevations in astrocytes might induce glutamate release (Santello et al., 2011). Moreover, a previous study has reported that 500 nM Aβ dimers increase extracellular glutamate concentrations (Zott et al., 2019). In this study, we found that the P2Y1 receptor pathway is involved in Aβ dimers-mediated fast-microdomain [Ca^2+^]_*i*_ elevations. It will be important to explore whether Aβ dimers actually enhance glutamate release from astrocytes and whether P2Y1 is involved in this pathway. This will be a key cellular signaling pathway in the very early stage of AD.

## 5. Data Availability Statement

All data needed to evaluate the conclusions in the study are present in the main text and the Supplementary Materials. Further study-related data could be requested from the authors.

## 6. Author Contributions

K.N. proposed the research project and all authors contributed to the conceptualization of the project. K.N. performed all experiments, K.O. contributed to the development of the optical imaging system, M.S. and J.S. contributed to the analysis and statistical evaluation of the data, and K.N., R.E., and T.N. contributed to the writing of the manuscript, and all authors approved the submitted version.

## 8. Acknowledgments

This work was supported in part by The Graduate University for Advanced Studies, SOKENDAI. We also thank the Genetically-Encoded Neuronal Indicator and Effector Project and the Janelia Farm Research Campus of the Howard Hughes Medical Institute for sharing the GCaMP6f constructs; Yuki Watakabe, Miwa Kawachi, Maki Watanabe, Miyoko Shimomura and Chiemi Hyodo for their help with animal care and laboratory management, and Kana Tsuchiya, Ayaka Osamura for excellent secretary works. We are also grateful to Profs. Toshiko Yamazawa (Jikei University), Hiroaki Wake (NIPS), Hideji Murakoshi (NIPS), and the members of Biophotonics Laboratory for helpful discussions of this study. The authors would like to thank enago (www.enago.jp) for the English language review.

## 10. Supplementary Material

