## Supplemental Material for "Amyloid-β-induced Alteration of Fast and Localized Calcium Elevations in Cultured Astrocytes"

### 1 Supplementary Data

**Supplementary Data Sheet 1. Summary of individual data and statistical results in the figures**

### 2 Supplementary Figures, Table, and Movies

#### Supplementary Figures

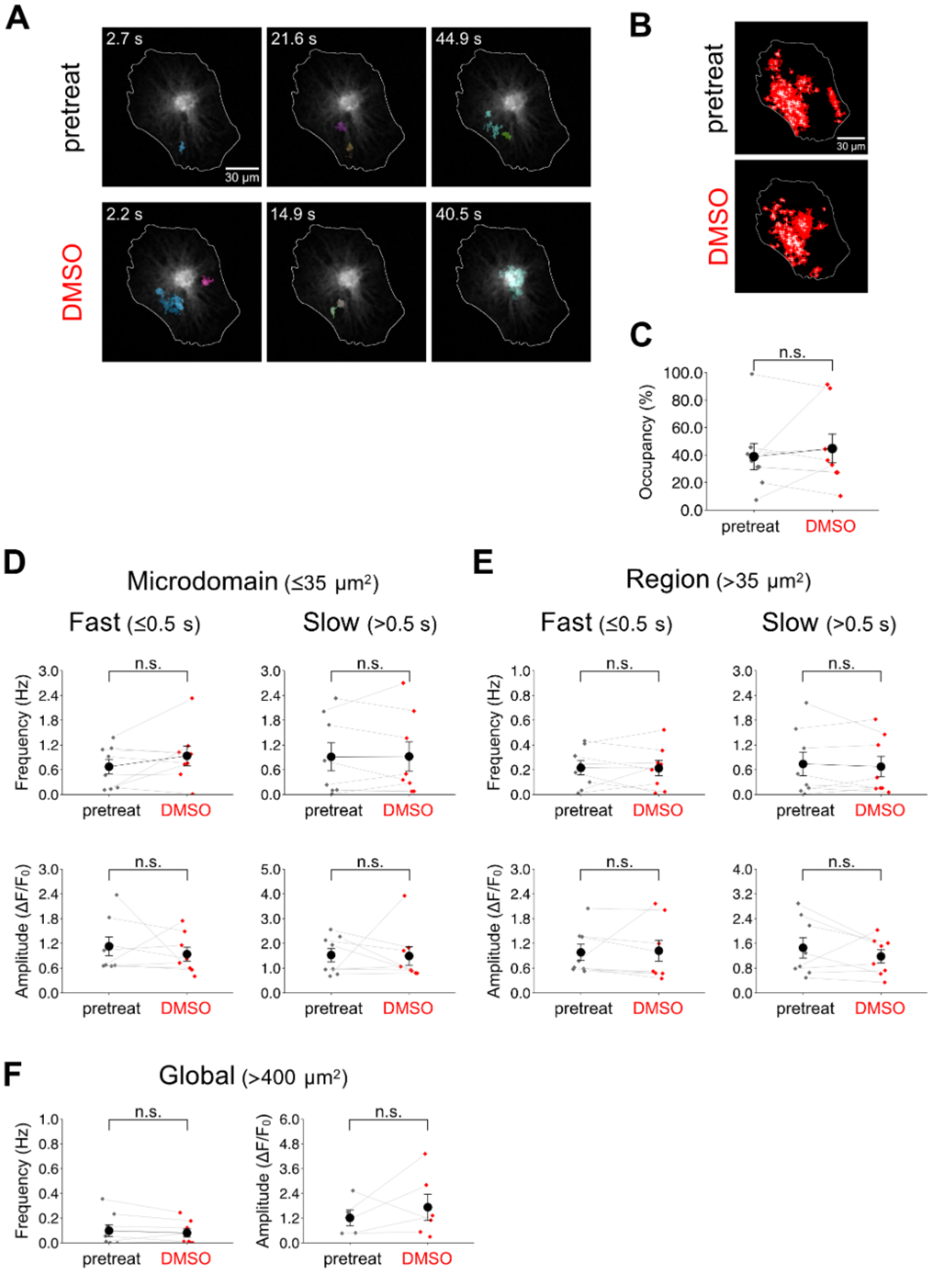


**Supplementary Figure 1. Effects of DMSO (1%) on $[Ca^{2+}]_i$ elevations in astrocytes**

**(A)** Representative images of astrocytic $[Ca^{2+}]_i$ signals are shown with AQuA-detected events. Images of pretreat (top) and under DMSO application (bottom) are shown. The time is shown in the upper left corner of each image. The total acquisition time is 90 s for each condition. **(B)** Comparison of the area of $[Ca^{2+}]_i$ elevations in each condition. The two images on the left represent the area of total of $[Ca^{2+}]_i$ elevations in the pretreat and DMSO application. The red area and the white cross indicate the area of $[Ca^{2+}]_i$ elevation and the center of gravity of individual $[Ca^{2+}]_i$ elevations, respectively. **(C)** Comparison of the event occupancy of $[Ca^{2+}]_i$ elevations between pretreat and DMSO application. **(D, E, F)** Frequencies (Hz) and amplitudes ($\Delta F/F_0$) of $[Ca^{2+}]_i$ elevations are compared between pretreat and DMSO application. Data are expressed as mean ± s.e.m. Two-sided Wilcoxon's signed rank sum test was used to compare differences (n = 8). Individual data and statistical results in the graphs are summarized in Supplemental Data Sheet 1. n.s.: not significant.

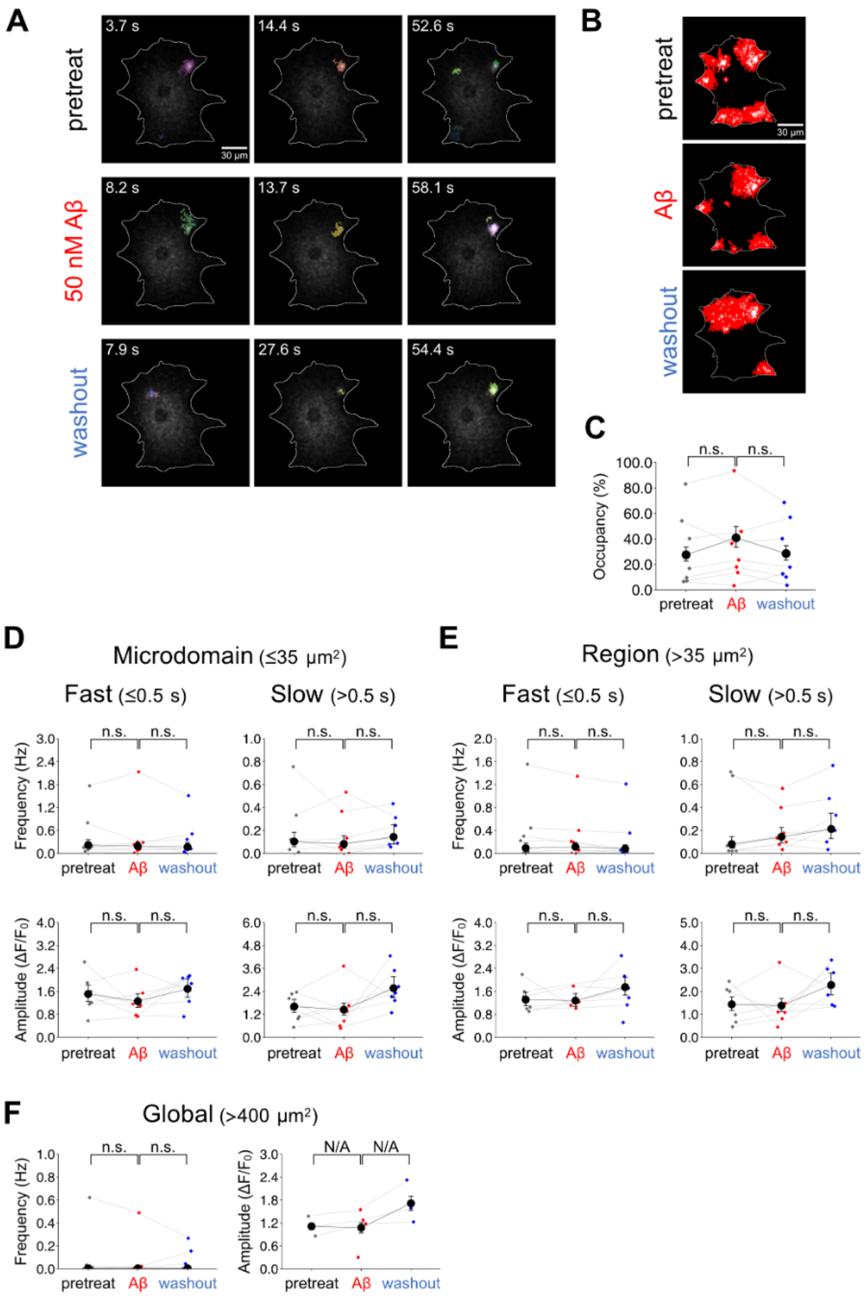


**Supplementary Figure 2. Effects of 50 nM Aβ dimers on $[Ca^{2+}]_i$ elevations in astrocytes**

See figure legend in Fig. S1 (n = 7). Data are expressed as estimate ± standard error. GLM or GLMM was used to evaluate the difference. n.s.: not significant, N/A: not applicable.

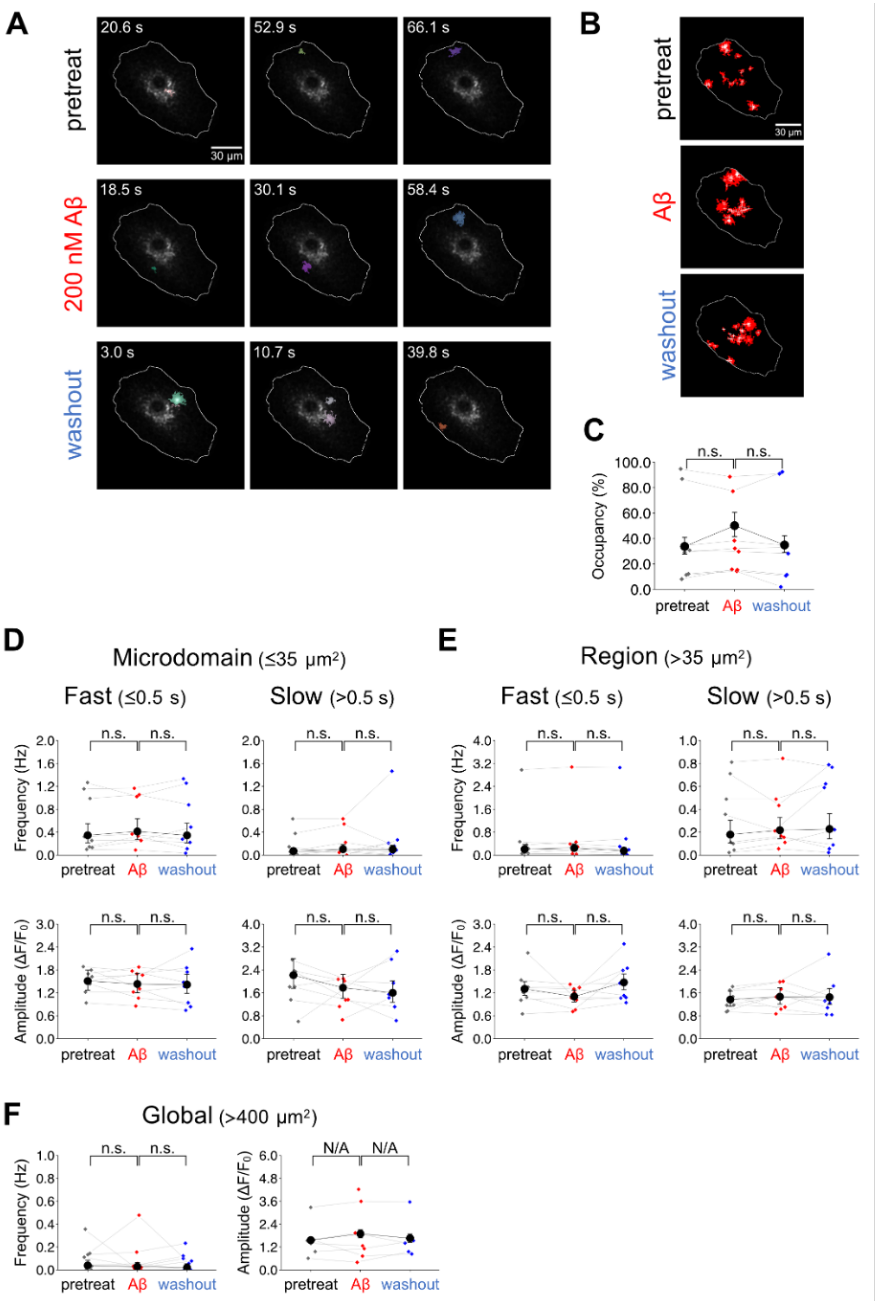


**Supplementary Figure 3. Effects of 200 nM Aβ dimers on $[Ca^{2+}]_i$ elevations in astrocytes**

See figure legend in Fig. S1 (n = 8). Data are expressed as estimate ± standard error. GLM or GLMM was used to evaluate the difference. n.s.: not significant, N/A: not applicable.

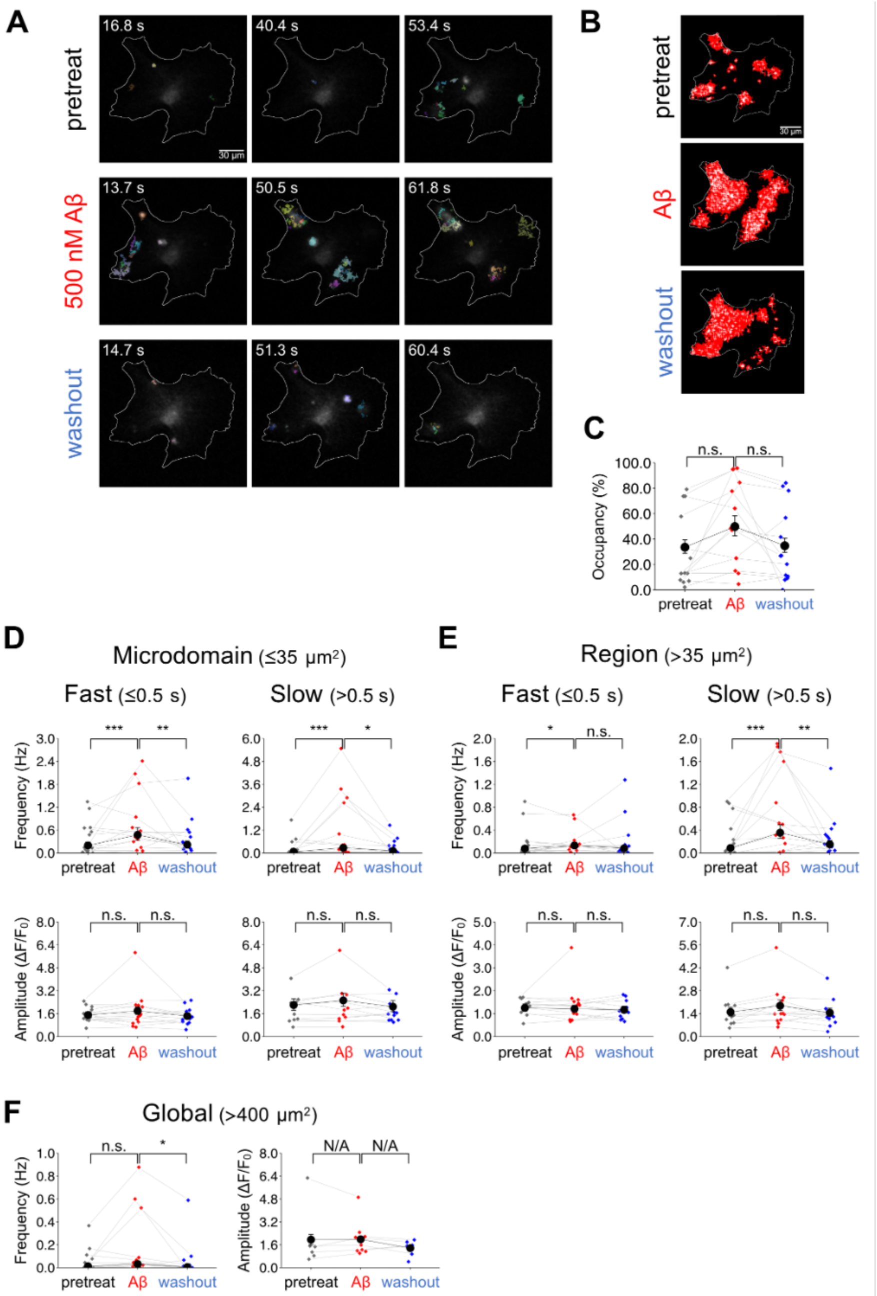


**Supplementary Figure 4. Effects of 500 nM Aβ dimers on $[Ca^{2+}]_i$ elevations in astrocytes**

See figure legend in Fig. S1 (n = 13). Data are expressed as estimate ± standard error. GLM or GLMM was used to evaluate the difference. n.s.: not significant, N/A: not applicable.

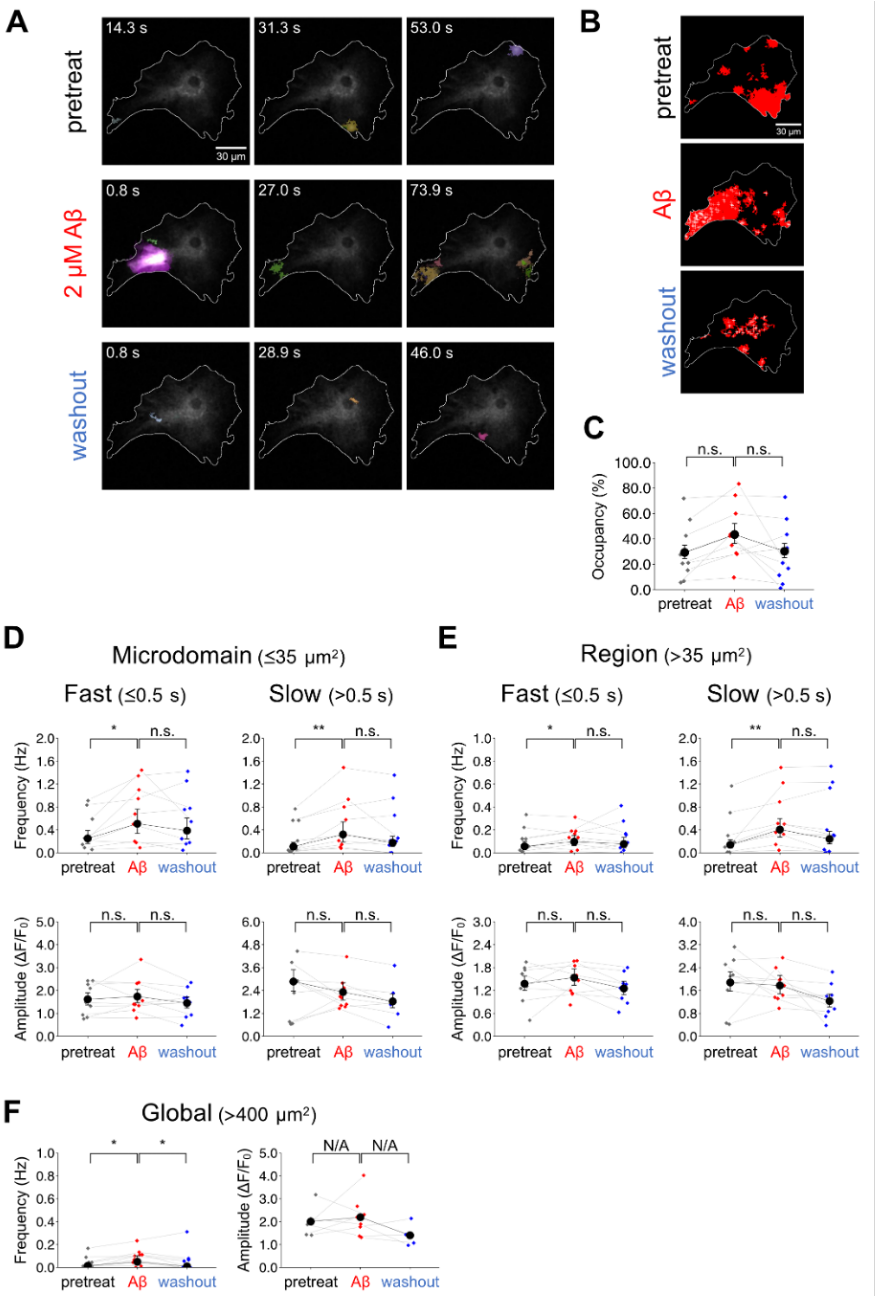


**Supplementary Figure 5. Effects of 2 µM Aβ dimers on $[Ca^{2+}]_i$ elevations in astrocytes**

See figure legend in Fig. S1 (n = 9). Data are expressed as estimate ± standard error. GLM or GLMM was used to evaluate the difference. n.s.: not significant, N/A: not applicable.

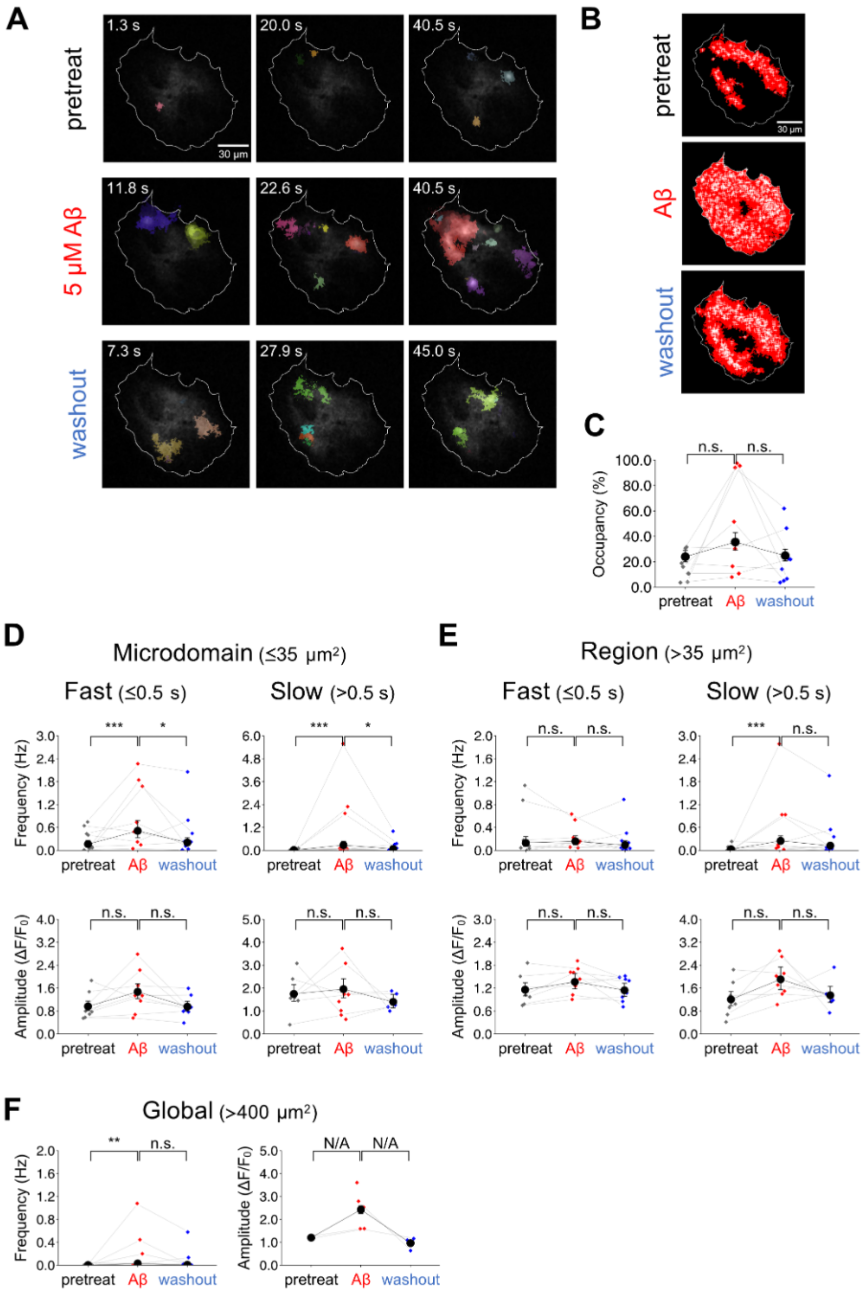


**Supplementary Figure 6. Effects of 5 µM Aβ dimers on $[Ca^{2+}]_i$ elevations in astrocytes**

See figure legend in Fig. S1 (n = 8). Data are expressed as estimate ± standard error. GLM or GLMM was used to evaluate the difference. n.s.: not significant, N/A: not applicable.

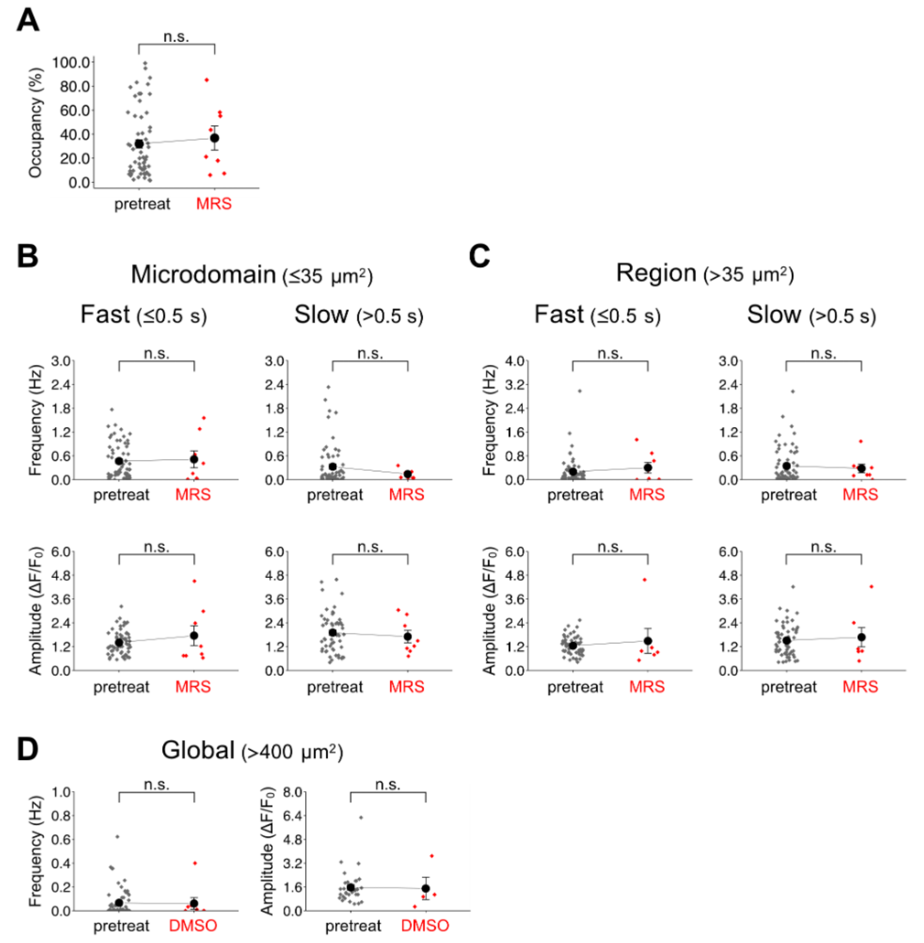


**Supplementary Figure 7. Effects of MRS2179 on [$Ca^{2+}$]$_i$ elevations in astrocytes**

See figure legend in Fig. S1 (pretreat (n = 70) vs MRS2179 (n = 8)). Two-sided Wilcoxon-Mann-Whitney test was used to evaluate the difference. n.s.: not significant.

**Supplementary Table**

**Supplementary Table 1. AQuA parameters used to detect $[Ca^{2+}]_i$ events in astrocytes**

| Parameters | Value |
|---|---|
| Signal: intensity threshold | 3 |
| Signal: smoothing | 0 |
| Signal: minimum size | 50 px |
| Voxel: temporal cut | 2 |
| Voxel: growing z | 1 |
| Event: rising time uncertainty | 2 |
| Event: slowest delay in propagation | 2 |
| Event: propagation smoothness | 1 |
| Clean: Z-score threshold | 3 |

**Supplementary Movie**

**Supplementary Movie 1. AQuA-detected $[Ca^{2+}]_i$ elevations before Aβ dimers applications.**

**Supplementary Movie 2. AQuA-detected $[Ca^{2+}]_i$ elevations under 500 nM Aβ dimers. applications**

**Supplementary Movie 3. AQuA-detected $[Ca^{2+}]_i$ elevations after washout.**
